# Carbon Assimilation Strategies in Ultrabasic Groundwater: Clues from the Integrated Study of a Serpentinization-Influenced Aquifer

**DOI:** 10.1101/776849

**Authors:** L.M. Seyler, W.J. Brazelton, C. McLean, L.I. Putman, A. Hyer, M.D.Y. Kubo, T. Hoehler, D. Cardace, M.O. Schrenk

## Abstract

Serpentinization is a low-temperature metamorphic process by which ultramafic rock chemically reacts with water. These reactions provide energy and materials that may be harnessed by chemosynthetic microbial communities at hydrothermal springs and in the subsurface. However, the biogeochemistry of microbial populations that inhabit these environments are understudied and are complicated by overlapping biotic and abiotic processes. We applied metagenomics, metatranscriptomics, and untargeted metabolomics techniques to environmental samples taken from the Coast Range Ophiolite Microbial Observatory (CROMO), a subsurface observatory consisting of twelve wells drilled into the ultramafic and serpentinite mélange of the Coast Range Ophiolite in California. Using a combination of DNA and RNA sequence data and mass spectrometry data, we determined that several carbon assimilation strategies, including the Calvin-Benson-Bassham cycle, the reverse tricarboxylic acid cycle, the reductive acetyl-CoA pathway, and methylotrophy are used by the microbial communities inhabiting the serpentinite-hosted aquifer. Our data also suggests that the microbial inhabitants of CROMO use products of the serpentinization process, including methane and formate, as carbon sources in a hyperalkaline environment where dissolved inorganic carbon is unavailable.

**Importance:** This study describes the metabolic pathways by which microbial communities in a serpentinite-influenced aquifer may produce biomass from the products of serpentinization. Serpentinization is a widespread geochemical process, taking place over large regions of the seafloor, particularly in slow-spreading mid ocean ridge and subduction zone environments. The serpentinization process is implicated in the origin of life on Earth and as a possible environment for the discovery of life on other worlds in our solar system. Because of the difficulty in delineating abiotic and biotic processes in these environments, major questions remain related to microbial contributions to the carbon cycle and physiological adaptation to serpentinite habitats. This research explores multiple mechanisms of carbon assimilation in serpentinite-hosted microbial communities.

## Introduction

Serpentinization is the process by which ultramafic rock in the lower crust and upper mantle of the Earth is hydrated, leading to the oxidation of ferrous iron in the minerals olivine and pyroxene, and the release of hydrogen (H_2_) and hydroxyl ions (OH^-^). The reducing power supplied by serpentinization, when mixed with oxidants from surface and subsurface fluids, creates chemical disequilibria that microbial populations can harness (McCollom, 2007; Amend et al., 2011; Cardace et al., 2015). However, due to the release of cations during rock weathering much of the dissolved inorganic carbon (DIC) in serpentinite systems is precipitated as calcite and aragonite (Barnes et al., 1978). Consequently, small carbon-bearing compounds including methane, carbon monoxide, and formate can take on important roles in sustaining microbial ecosystems in serpentinites. The high pH and limited availability of both DIC and electron acceptors creates a challenging environment for microbial communities hosted in serpentinization-influenced environments (Twing et al. 2017).

Overlapping biogenic and abiogenic processes in serpentinizing rocks make it difficult to delineate carbon cycling pathways used in these environments (Figure 1). Under highly reducing conditions, in the presence of specific mineral catalysts, hydrogen reacts with carbon dioxide (CO_2_) or carbon monoxide (CO) to form methane (CH_4_) and small-chain hydrocarbons via Fischer-Tropsch type reactions (McCollom and Seewald, 2001; Charlou et al., 2002; Proskurowski et al., 2008; McCollom, 2013). These same compounds may also be formed by biological activity, or diagenesis of organic matter (Sephton and Hazen, 2013). For example, high concentrations of dissolved organic material, produced by past microbiological activity that may have been supported by the by-products of serpentinization, have been detected in close association with serpentine-hosted hydrogarnets recovered from the Mid-Atlantic Ridge (Ménez et al., 2012). Acetate in endmember fluids at Lost City is also likely produced by biological activity (Lang et al., 2010), but other hydrocarbons in the rock-hosted fluids do not have a clear biotic or abiotic source (Delacour et al., 2008), though formate and C_2_^+^ alkanes appear to be produced abiotically (Proskurowski et al., 2008; Lang et al., 2010). Biotic versus abiotic sources of methane in serpentinizing systems are likewise difficult to resolve and vary from site to site (Bradley and Summons, 2010; Etiope et al., 2013; Wang et al., 2015; Kohl et al., 2016).

**Figure 1.**
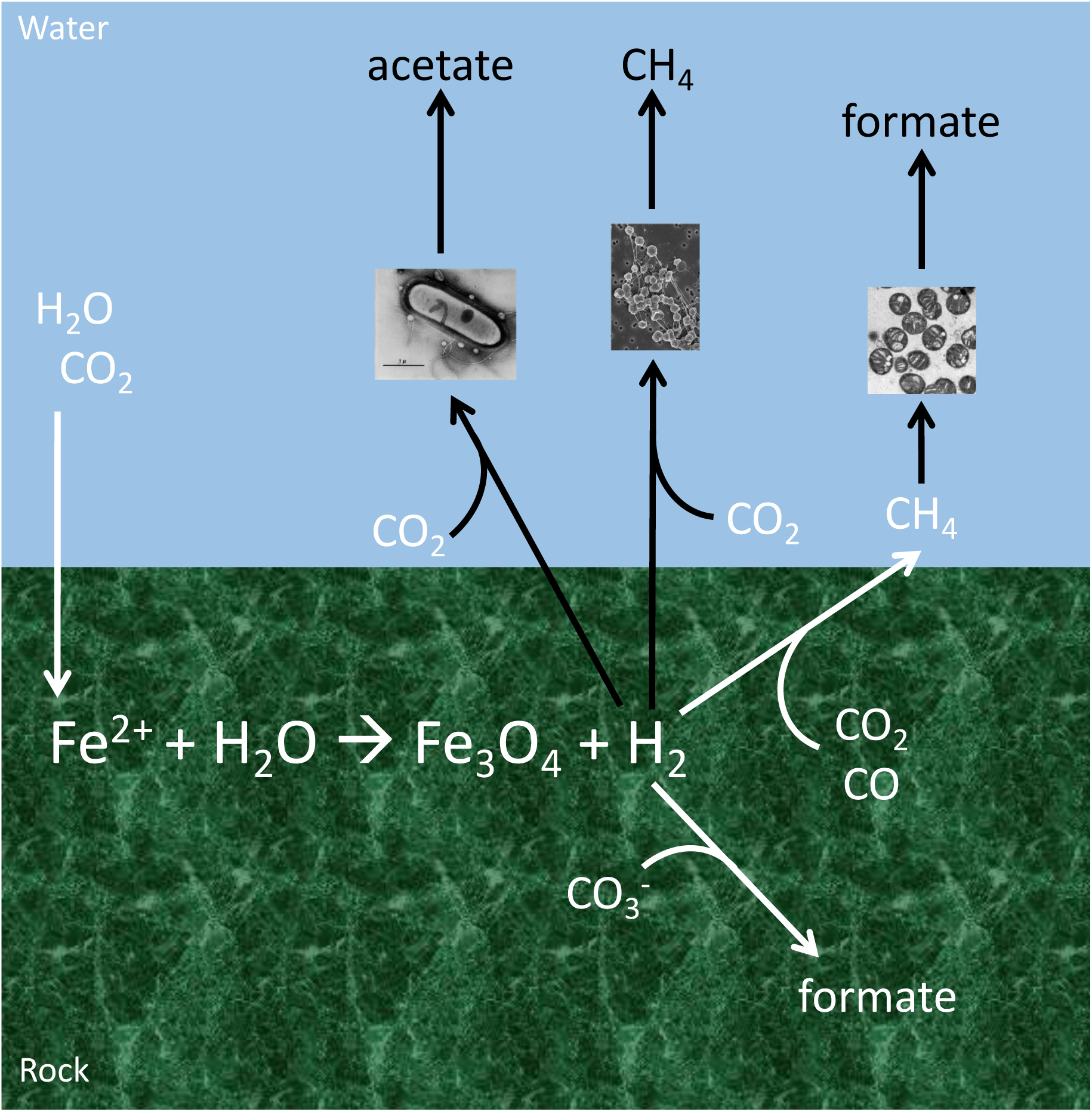
A diagram depicting abiotic (white) and biotic (black) sources of carbon compounds in serpentinizing environments. Adapted from Preiner et al., 2018.

*In situ* microbial communities may utilize the products of serpentinization - including methane, carbon monoxide, formate, and acetate - as sources of carbon (Schrenk et al., 2004; Lang et al., 2010; Brazelton et al., 2011; Brazelton et al., 2012; Lang et al., 2012; Brazelton et al., 2013; Crespo-Medina et al., 2014). Microbial communities in serpentinite springs in the Voltri Massif, Italy, appear to perform both aerobic methanotrophy and methanogenesis using CO_2_ liberated from acetate (Brazelton et al., 2017). Methanogens in The Cedars, CA, are capable of producing methane from both inorganic carbon and a variety of organic carbon substrates: acetate may be produced through fermentation (Kohl et al., 2016), and communities that reside deeper in the springs also possess a reductive acetyl-CoA pathway for autotrophic acetogenesis (Suzuki et al., 2017). Carbon monoxide may also be used as a substrate by microbial communities in the Tablelands, Canada, but it appears to be primarily used as a source of energy rather than carbon (Morrill et al., 2014). The carbonate chimneys of the Lost City Hydrothermal Field are dominated by a single species of archaea that may be capable of both production and consumption of methane in tandem (Brazelton et al., 2011). Genes encoding NAD(+)-dependent formate dehydrogenase in metagenomes recovered from the Samail Ophiolite aquifer suggest that formate is an important source of carbon in these DIC-limited hyperalkaline waters (Fones et al., 2019). In short, carbon assimilation in these environments is both enigmatic to researchers and challenging for in situ microbial communities, but pathways and strategies utilizing alternate sources of carbon supplied by serpentinization appear to be prevalent.

In this study, we examined the microbial metabolic potential and biogeochemistry of a serpentinization-influenced aquifer in the Coast Range Ophiolite Microbial Observatory (CROMO), California. Wells have been drilled at various depths and locations at CROMO such that microbial processes can be explored across a wide range of physical and chemical conditions. Carbon assimilation pathways in the microbial communities at CROMO were investigated using a combination of metagenomic, metatranscriptomic, and metabolomic approaches. Metagenomic data sets from nine of the twelve CROMO wells, and metatranscriptomic data sets from four wells, obtained over multiple sampling trips, were queried for complete pathways and individual genes associated with carbon assimilation. We then applied environmental metabolomics, an emerging approach used to directly assess the products of ecosystem activity, on intra- and extracellular extracts from the two most alkaline wells (CSW1.1 and QV1.1) to describe the dissolved organic carbon pools in these wells. This study links the genetic potential of serpentinite-hosted communities to the observed biogeochemistry of the groundwater in CROMO.

## Materials and Methods

### Site Description

The Coast Range Ophiolite in northern California is a 155–170 million-year-old portion of oceanic crust tectonically emplaced on the North American continental margin. Trapped Cretaceous seawater and circulating meteoric water reacts with ultramafic rock (Peters, 1993; Shervais et al., 2005) as evidenced by numerous calcium hydroxide rich springs throughout the formation (Barnes and O’Neil, 1971). Previous studies of continental serpentinite environments have sampled similar springs in the Tablelands, Newfoundland, Canada (Brazelton et al., 2012; Brazelton et al., 2013) and The Cedars, California (Suzuki et al., 2013), providing information on the surface expression of subsurface biogeochemical processes. In order to directly access the serpentinite subsurface environment and monitor groundwater biogeochemistry with minimal contamination, a microbial observatory was established in the Coast Range Ophiolite at the UC-Davis McLaughlin Natural Reserve (Lower Lake, CA) (Cardace et al., 2013). The Coast Range Ophiolite Microbial Observatory (CROMO) consists of two sets of wells 1.4 km apart—the Quarry Valley (QV) and Core Shed Wells (CSW)—each with an uncased main borehole (designated “1.1”) surrounded by two or four monitoring wells, respectively. In addition to the eight newly drilled wells, monitoring wells (N08A, N08B, and N08C at Quarry Valley, and CSWold) were previously drilled by the Homestake Mining Company, Inc. more than 30 years prior, in conjunction with mining operations conducted in the area. All of the wells are cased with PVC, with the exception of the 1.1 wells which are only cased to bedrock, leaving the bottom of the hole uncased.

### Extraction of DNA and RNA

For each environmental sample from the CROMO wells, four liters of fluid were pumped via slow flow bladder pump (Cardace et al., 2013) and immediately filtered through Sterivex 0.2 µm filter cartridges (Millipore, Billerica, MA) using a portable peristaltic pump (Masterflex®, Cole Parmer, Vernon Hills, IL). Cartridges were kept on ice during filtration, immediately stored in liquid nitrogen upon completion, shipped to the home laboratory, and stored at -80°C until processing. Total genomic DNA extractions were completed as previously described by Brazelton et al., (2017), Crespo-Medina et al., (2017) and Twing et al., (2017) and briefly described here. Freeze/thaw cycles and lysozyme/Proteinase K treatment were performed to lyse cells, followed by purification with phenol-chloroform, precipitation using ethanol, and purification using QiaAmp (Qiagen, Hilden, Germany) columns according to the manufacturer’s instructions. A Qubit 2.0 fluorometer (ThermoFisher, Waltham, MA) was used to quantify extracted DNA using a Qubit dsDNA High Sensitivity Assay kit.

Extractions for RNA from CROMO wells were performed as described previously with slight modifications (Lin and Stahl, 1995; MacGregor et al., 1997). Briefly, frozen 0.2 µm Sterivex filter cartridges were broken open, cut into four equal pieces, and divided into two screw-cap Eppendorf tubes containing phenol, 20% sodium dodecyl sulfate, 5× low-pH buffer, and 0.2 to 0.5 g baked zirconium beads. Samples were bead-beaten for 3 minutes, heated in a 60°C water bath for 10 minutes, bead beaten again for 3 minutes, and centrifuged at 4°C and 18,407 × *g* to separate phases. Supernatant was transferred to a fresh Eppendorf tube and chilled. 1× low-pH buffer was added to the remaining sample, and bead-beating was repeated. Supernatants were combined, and phenol, 1:1 phenol:chloroform, and chloroform were added in series with vortexing and centrifugation in between. Between steps, aqueous phases were transferred to clean Eppendorf tubes. The final aqueous phase was transferred to a clean Eppendorf tube with additions of ammonium acetate, isopropanol, and magnesium chloride before vortexing and incubation at -20°C overnight. Samples were centrifuged for 30 minutes at 4°C, washed with ethanol, and dried under vacuum before suspension in RNase-free water and storage at -80°C until analyzed.

### Metagenome and Metatranscriptome Analyses

This study reports a new analysis of the metagenome and metatranscriptome sequences reported in Sabuda et al. (in review). Our metagenomic assembly and mapping of metatranscriptome reads was identical to that of Sabuda et al., and all methods are reported again here for completeness. Metagenome and metatranscriptome sequencing was conducted at the Joint Genome Institute (JGI) on an Illumina HiSeq2000 instrument. A Covaris LE220 ultrasonicator was used to shear DNA samples into 270 bp fragments, and size selection was performed using solid phase reversible immobilization (SPRI) magnetic beads (Rohland and Reich, 2012). DNA fragments were end-repaired, A-tailed, and ligated with Illumina-compatible adapters with barcodes unique for each library. KAPA Biosystem’s next-generation sequencing library qPCR kit and Roche LightCycler 280 RT PCR instrument were used to quantify libraries. 10-library pools were assembled and prepared for Illumina sequencing in one lane each.

Clustered flowcells were produced using a TruSeq paired-end cluster kit (v3) and Illumina’s cBot instrument. The Illumina HiSeq2000 instrument was utilized with a TruSeq SBS sequencing kit (v3) and a 2 × 150 indexed run recipe to sequence the samples. The raw sequence reads were trimmed by the JGI with a minimum quality score cutoff of 10 and to remove adapters. These trimmed reads from CROMO wells were previously reported by Twing et al., 2017, but additional quality-filtering and a new assembly, distinct from the JGI assembly reported by Twing et al., was performed for this study.

The trimmed reads from the JGI were subjected to an additional quality screen to trim 3’ adapters with cutadapt v. 1.15 (Martin, 2011), to remove replicate sequences, and to trim sequences again with a threshold of 20 along a sliding window of 6 bases with qtrim (part of the seq-qc package: https://github.com/Brazelton-Lab/seq-qc). All CROMO metagenomes and metatranscriptomes were pooled together for a master CROMO assembly computed with Ray Meta v.2.3.1 (Boisvert et al., 2012). Phylogenetic affiliation of contigs was assigned using PhyloPythiaS+ (Gregor et al., 2016). Metagenome and metatranscriptome short reads were mapped to the pooled assembly using Bowtie2 v.2.2.6 (Langmead and Salzberg, 2013). The Prokka pipeline (Seemann, 2014) was used for gene prediction and functional annotation of contigs. The arguments –metagenome and –proteins were used with Prokka v.1.12 to indicate that genes should be predicted with the implementation of Prodigal v.2.6.2 (Hyatt et al., 2010) optimized for metagenomes as described by Twing et al., 2017. Metagenome-assembled genome (MAG) bins were constructed with ABAWACA (https://github.com/CK7/abawaca) using tetranucleotide frequencies and differential abundance as measured by Bowtie2 mapped read abundances. Bin quality was computed with CheckM (Parks et al., 2015), and only “high quality” MAG bins are reported here (>50% completeness, <10% contamination, as defined by Bowers et al., 2017). The completeness and contamination of some bins were improved by refinement with Binsanity (Graham et al., 2017). Taxonomic classifications of MAGs were annotated using GTDB-Tk (https://github.com/Ecogenomics/GTDBTk; Parks et al., 2018).

Predicted protein-coding sequences were annotated by searching the Kyoto Encyclopedia of Genes and Genomes (KEGG; Ogata et al., 1999; Kanehisa and Goto, 2000) release v. 83.2 within Prokka. HTSeq v.0.6.1 (Anders et al., 2015) and were used to calculate predicted protein abundances. The abundances of predicted protein functions in all CROMO metagenomes and metatranscriptomes were normalized to predicted protein size and metagenome size. Data reported here are in units of metagenome fragments per kilobase of predicted protein sequence per million mapped reads (FPKM). Abundances of metabolic pathways were obtained by mapping KEGG protein IDs and their normalized counts onto the FOAM ontology (Prestat et al., 2014) with MinPath (Ye & Doak, 2009) as implemented in HUMAnN2 v.0.6.0 (Abubucker et al., 2012).

The complete translated genome assemblies of Serpentinomonas isolates from the Cedars (Comamonadaceae bacterium A1, B1, and H1) were obtained as amino acid FASTA files from RefSeq (accession numbers: GCF_000828895.1, GCF_000828915.1, GCF_000696225.1). ORFs were annotated as KEGG orthologs (KOs) using BlastKOALA (Kanehisa and Goto 2000). KEGG modules were annotated within genomes using the KEGG module reconstruction tool. MAGs were translated to protein sequences using Prodigal (Hyatt et al., 2010) and annotated in the same manner. Modules within the three cultured representative genomes and the MAGs were then directly compared in order to compare pathways identified in available cultivars from the Cedars to pathways in the uncultured communities hosted in CROMO’s hyperalkaline groundwater. A phylogenetic tree comparing MAGs to the *Serpentinomonas* isolates was generated using SpeciesTree in KBase (Arkin et al., 2018).

### Metabolite Extraction and Analysis

Three 4-L samples of groundwater from QV1.1 and CSW1.1 were taken in June 2016 and filtered using 0.22 µm 47 mm polytetrafluoroethylene (PTFE) filters on a glass vacuum filtration tower. Filters were frozen in liquid nitrogen and stored at -80° C until extraction (Fiore et al., 2015; Kido Soule et al., 2015). Approximately 30 mL filtrate was set aside and frozen at - 20 °C for total organic carbon analysis on a Shimadzu Total Organic Carbon Analyzer. Formaldehyde was measured using a CHEMetrics™ Vacu-vials™ colorimetric kit (detection range 0.4-8.0 ppm). The remaining filtrate was acidified to pH 2-3 using concentrated HCl and stored at 4°C for 5 days until further analysis by solid phase extraction.

Frozen filters were cut into small pieces over a muffled piece of aluminum foil, using methanol-washed scissors, then transferred into muffled amber glass vials. 2 mL cold extraction solvent (40:40:20 acetonitrile:methanol:0.1 M formic acid) was added to each vial, and the samples were sonicated for 10 minutes. Extracts were transferred to a centrifuge tube via pipette and centrifuged at 20,000 × g for 5 min at 4 °C. The supernatant was then transferred to a new amber glass vial, neutralized with 6 M ammonium hydroxide, and vacuum centrifuged to dryness. Samples were dissolved in 495 µl 95:5 water:acetonitrile with 5 µl of 5 µg/ml of biotin-(ring-6,6-d_2_) added as an injection standard (Rabinowitz and Kimball, 2007; Fiore et al., 2015).

Dissolved organics were captured from the acidified filtrate on solid phase extraction (SPE) Bond Elut-PPL cartridges (Agilent Technologies) and eluted with 100% methanol into muffled amber glass vials (Dittmar et al., 2008; Fiore et al., 2015). SPE-PPL cartridges were rinsed with methanol immediately before use. The supernatant was passed through the cartridge using 1/8” × 1/4” PTFE tubing to pull the supernatant into the cartridge to minimize the possibility of contamination from plastic leaching, while Viton tubing was used to remove the discarded flow-through via peristaltic pump (flow rate not exceeding 40 ml/min). The cartridges were then rinsed with at least 2 cartridge volumes of 0.01 M HCl to remove salt, and the sorbent dried with air for 5 minutes. Dissolved Organic Matter (DOM) was eluted from the cartridge with 2 ml methanol via gravity flow into a muffled amber glass vial. The eluate was then vacuum centrifuged to dryness. Precipitated samples were dissolved in 495 µl 95:5 water:acetonitrile, with 5 µl of 5 µg/ml of biotin-(ring-6,6-d_2_).

The above protocol was repeated for 4 L milliQ water, and organics extracted from the filter and filtrate as above as extraction blanks. 495 µl 95:5 water:acetonitrile plus 5 µl 5 µg/mL biotin-(ring-6,6-d_2_) was also run as a blank.

Samples were analyzed using tandem liquid chromatography mass spectrometry (LC/MS/MS) at the Michigan State University Metabolomics Core facility. Triplicate samples from each well were separated chromatographically in a Acquity UPLC BEH C18 column (1.7 µm, 2.1 mm × 50 mm) using a polar/non-polar gradient made up of 10 mM TBA and 15 mM acetic acid in 97:3 water:methanol (solvent A), and 100% methanol (solvent B). The gradient was run at 99.9/0.1 (% A/B) to 1.0/99.0 (% A/B) over 9 min, held an additional 3 min at 1.0/99.0 (% A/B), then reversed to 99.9/0.1 (% A/B) and held another 3 min. At a rate of 400µL/min, the sample was fed into a quadrupole time-of-flight mass spectrometer using a Waters Xevo G2-XS MS/MS in negative ion mode using a data independent collection method (m/z acquisition range 50 to 1500 Da). OM levels were too low to allow distinction between samples and blanks from the first run, so triplicate samples were pooled and run as before. After data was converted to centroid mode, raw data files were converted from proprietary Waters RAW format into XML-based mzML format using MSConvert (Chambers et al, 2012). Data was processed using XCMS (Smith et al., 2006) parametrized by AutoTuner (McLean and Kujawinski, in review). Isotopes and adducts of each feature were identified using CAMERA (Kuhl et al., 2012). This approach yielded a table of peak areas that are identified by a unique mass to charge (m/z) and retention time (rt) pair referred to as features. The table of features was subject to quality assurance through blank correction by removing any feature from a sample that also appeared within the blank control (described above) (Broadhurst et al., 2018). Features were first putatively annotated as KEGG compounds using mummichog (Li et al., 2013). Chemical standards of putative annotations were purchased whenever available to check the retention time to increase confidence of annotation from level 1 to 2 (Sumner et al. 2007). Due to the great complexity of the secondary mass spectral data and insufficient coverage of mass spectral databases for natural organic matter, MS2 information could not be readily used to increase confidence of annotation.

### Data Availability

CROMO metagenome and metatranscriptome sequences are publicly available in the NCBI SRA with the BioProject IDs PRJNA410019, PRJNA410020, PRJNA410022-410033, PRJNA410035-410037, PRJNA410054, PRJNA410057, PRJNA410286, PRJNA410403, PRJNA410404, PRJNA410553-410555, PRJNA410557. Raw spectrometry data are available in the MetaboLights database under study identifier MTBLS1260.

## Results

### Presence and Expression of Carbon Assimilation Pathways

The metagenomic data collected from nine of the twelve CROMO wells contains evidence for a range of carbon fixation/assimilation pathways (Figure 2). Functional profiling of the metagenomes using HUMAnN2 identified multiple C1 compound utilization and assimilation pathways, including the reductive acetyl-CoA (Wood-Ljungdahl) pathway, the reverse tricarboxylic acid (TCA) cycle, formaldehyde oxidation to formate, and formaldehyde fixation to biomass. Whenever these complete pathways were identified in the metagenomic data, they were also detected in the corresponding metatranscriptomic data (Figure 2). A complete Calvin-Benson-Bassham cycle was also detected in all nine metagenomes and all four metatranscriptomes, at a similar coverage across wells (Supplemental Figure 1). Carbon fixation pathways found exclusively in archaea (i.e., hydroxypropionate-hydroxybutylate cycle, 3-hydroxypropionate bicycle) were not detected. MAG bins contained the reductive pentose phosphate/Calvin-Benson-Bassham cycle, reductive TCA cycle, and reductive acetyl-CoA pathways (Figure 3).

**Figure 2.**
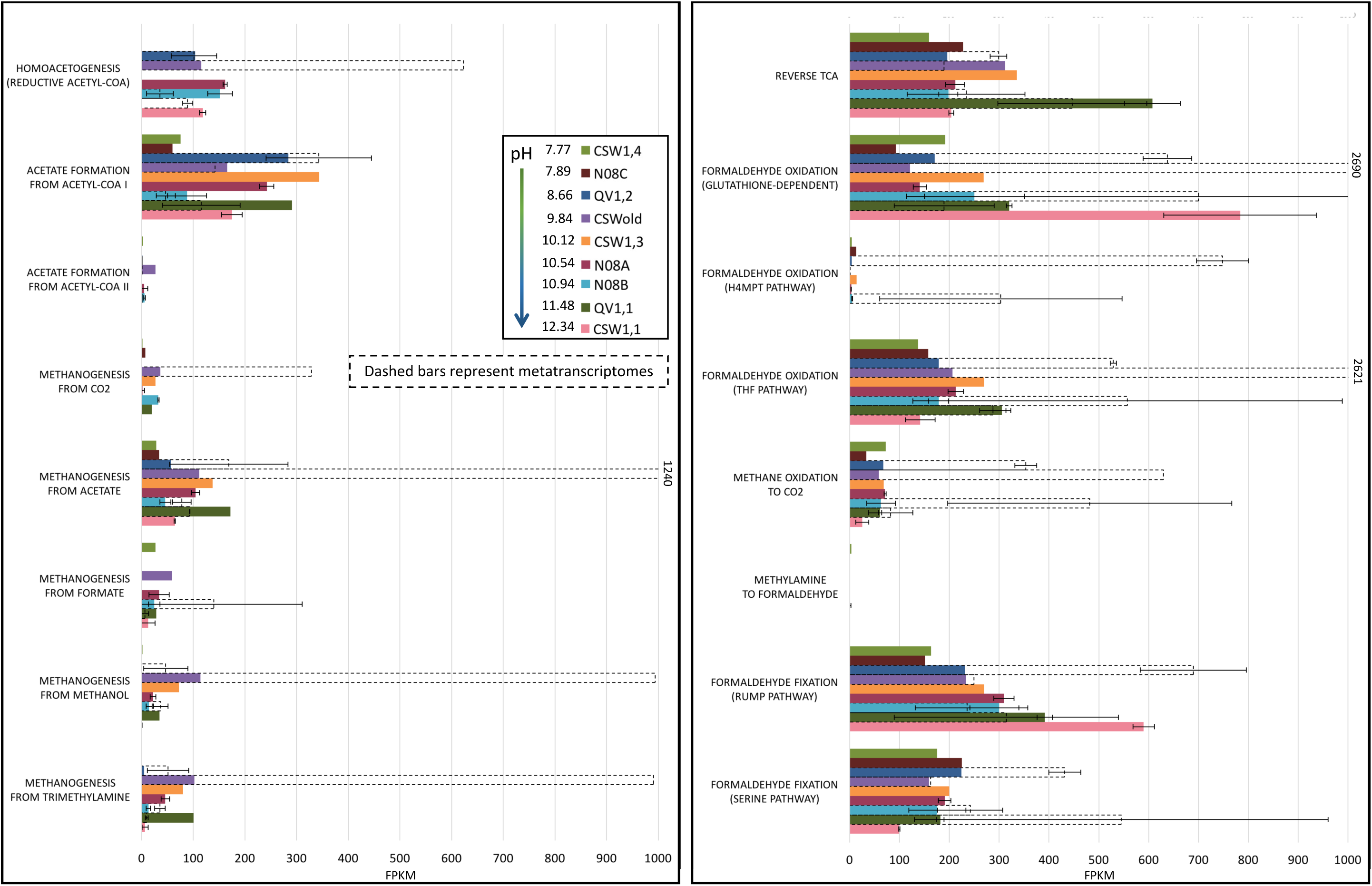
Metagenome coverage of Functional Ontology Assignments for Metagenomes (FOAM) pathways assigned using HUMAnN2. If a well was sampled for metagenomics more than once, the average percent coverage and standard error of the mean for that well are provided. Metatranscriptome coverage for four wells sampled for RNA is indicated using bars with dotted outlines.

**Figure 3.**
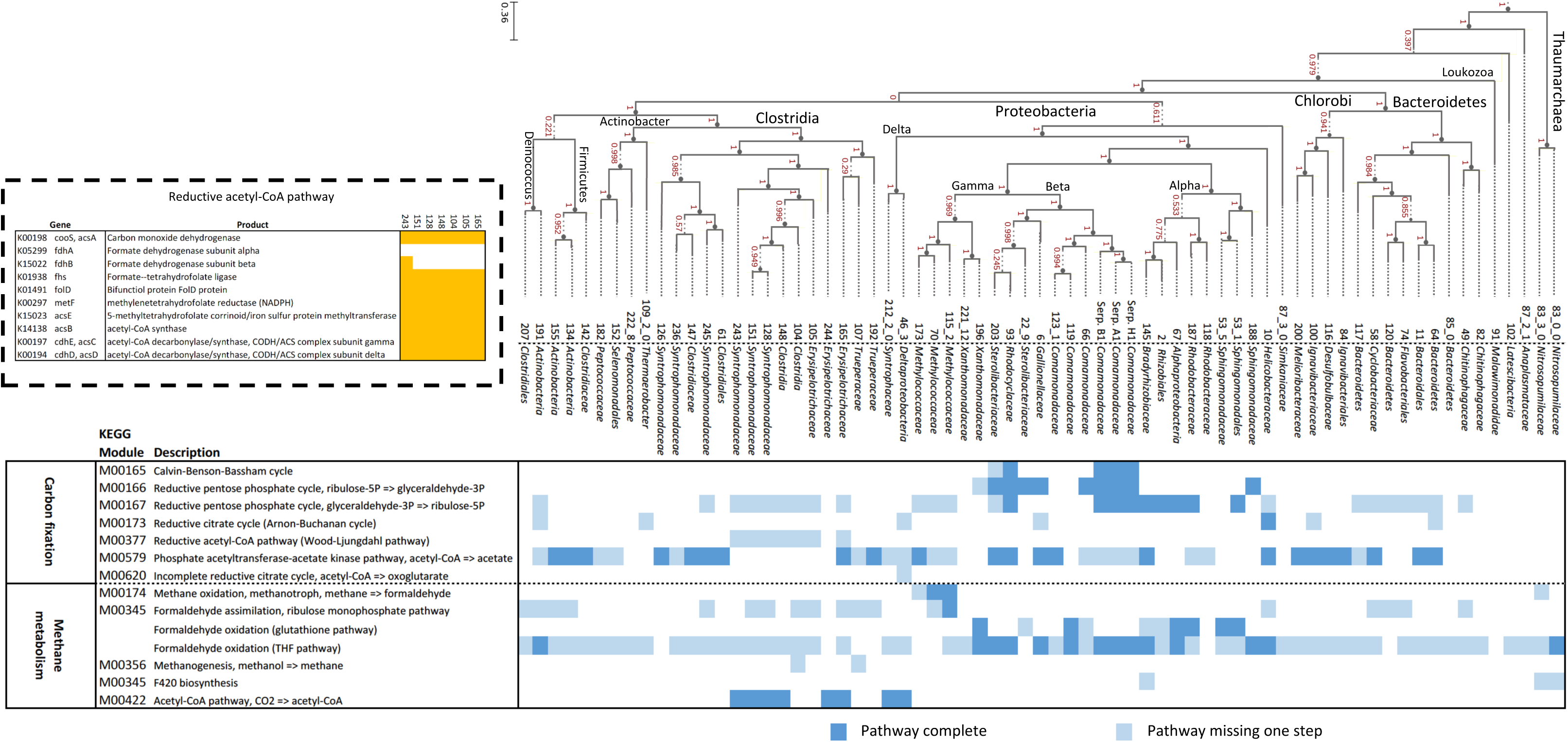
Selection of carbon cycling-associated KEGG modules detected in MAG bins, and *Serpentinomonas* isolate genomes (Suzuki et al., 2014), using BLASTKoala. Dark blue boxes indicate the complete module is present; light blue boxes indicate one step of the pathway is missing. The phylogenetic tree comparing the MAGs to the three *Serpentimonas* isolates was generated using SpeciesTree in KBase. Inset: Bins containing reductive acetyl-CoA pathway; gene presence in the genome is indicated in yellow.

Several pathways associated with methanotrophy and methylotrophy were detected in the metagenomes and metatranscriptomes. The pathway for aerobic methane oxidation to formaldehyde was present in most of the metagenomes (Figure 4) and the complete pathway was actively transcribed in wells QV1.2 and N08B. Formaldehyde uptake and assimilation pathways were prevalent in all of the sequenced wells (Figure 2). Complete glutathione-dependent and tetrahydrofolate (THF) pathways for formaldehyde oxidation to formate were present in all of the metagenomes, expressed in all four metatranscriptomes (Figure 2, Figure 4), and present in multiple MAGs (Figure 3). (The *gfa* gene, which codes for an enzyme that catalyzes a thermodynamically spontaneous step in the glutathione-dependent pathway [Goenrich et al., 2002], is only found in some formaldehyde-oxidizing bacteria and was absent from most of the metagenomes.) Formaldehyde fixation via the tetrahydromethanopterin (H4MPT) pathway was poorly represented across the metagenomes, but was expressed in the metatranscriptomes of QV1.2 and N08B. The capability for formaldehyde fixation to biomass by the ribulose monophosphate (RuMP) and serine pathways was likewise present in all nine metagenomes and all four metatranscriptomes (Figure 2), and the RuMP pathway was partially complete in 18 MAGs and complete in one MAG (Figure 3).

**Figure 4.**
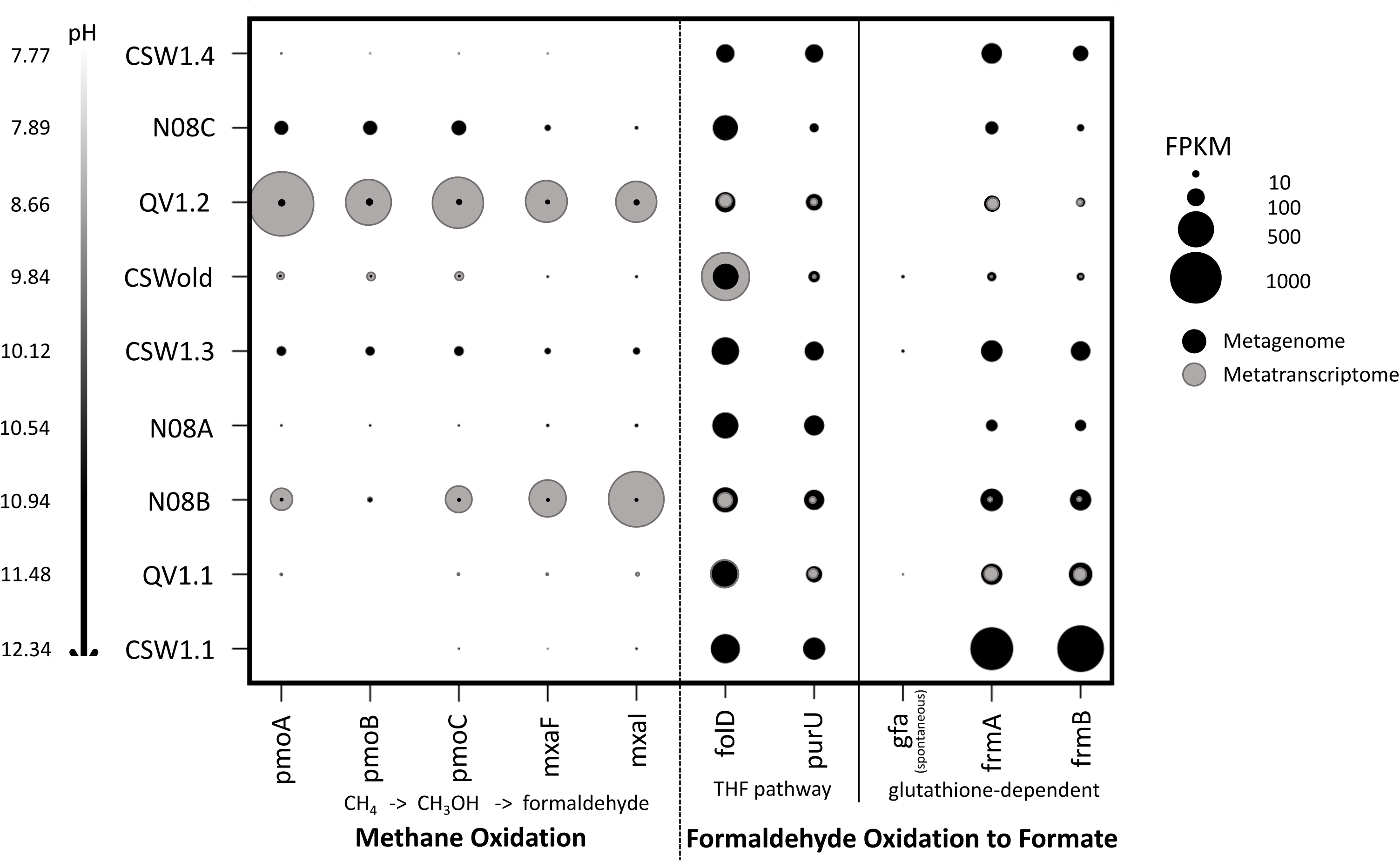
Fragments per kilobase of sequences per million mapped reads (FPKM) of genes in the homoacetogenic reductive acetyl-CoA pathway. Metagenomes are represented in black; metatranscriptomes are represented in gray. The genes for the two subunits of formate dehydrogenase (fdhA and fdhB) are missing or comparatively low in most wells. This enzyme is responsible for converting CO_2_ to formate.

Pathway identification using the KEGG database and FOAM pathways suggested that homoacetogenesis via the reductive acetyl-CoA pathway (also known as the Wood-Ljungdahl pathway) was represented in the metagenomes of only half of the wells sequenced (N08A, N08B, QV1.2, CSW1.1, and CSWold) (Figure 2). However, a gene-by-gene search indicated that the pathway was also present and nearly complete in the QV1.1 and CSW1.3 wells (Figure 5; Supplemental Tables 1 and 2). In the three least alkaline wells (CSW1.4, N08C, and QV1.2), most of the genes in the pathway were poorly represented (< 2 FPKM) in the metagenomic data. Metatranscriptomic data showed that the pathway was actively transcribed in all of the four wells that were sampled for mRNA (Figure 2; Figure 5). However, one of the initial steps of the reductive acetyl-CoA pathway, the conversion of CO_2_ to formate by NADP-dependent formate dehydrogenase (FDH), appeared to be poorly expressed or altogether missing in both the metagenomes and the metatranscriptomes (Figure 5). Seven of the 66 MAG bins also contained a nearly complete reductive acetyl-CoA pathway that was missing the genes encoding FDH (Figure 3, inset). The type I pathway for the formation of acetate from acetyl-CoA via acetyl-phosphate (present in Methanosarcina and Clostridiales) is relatively abundant across all metagenomes, while the type II pathway (which does not utilize an acetyl-phosphate intermediate) is sparsely represented in the metagenomic data (Figure 2).

**Figure 5.**
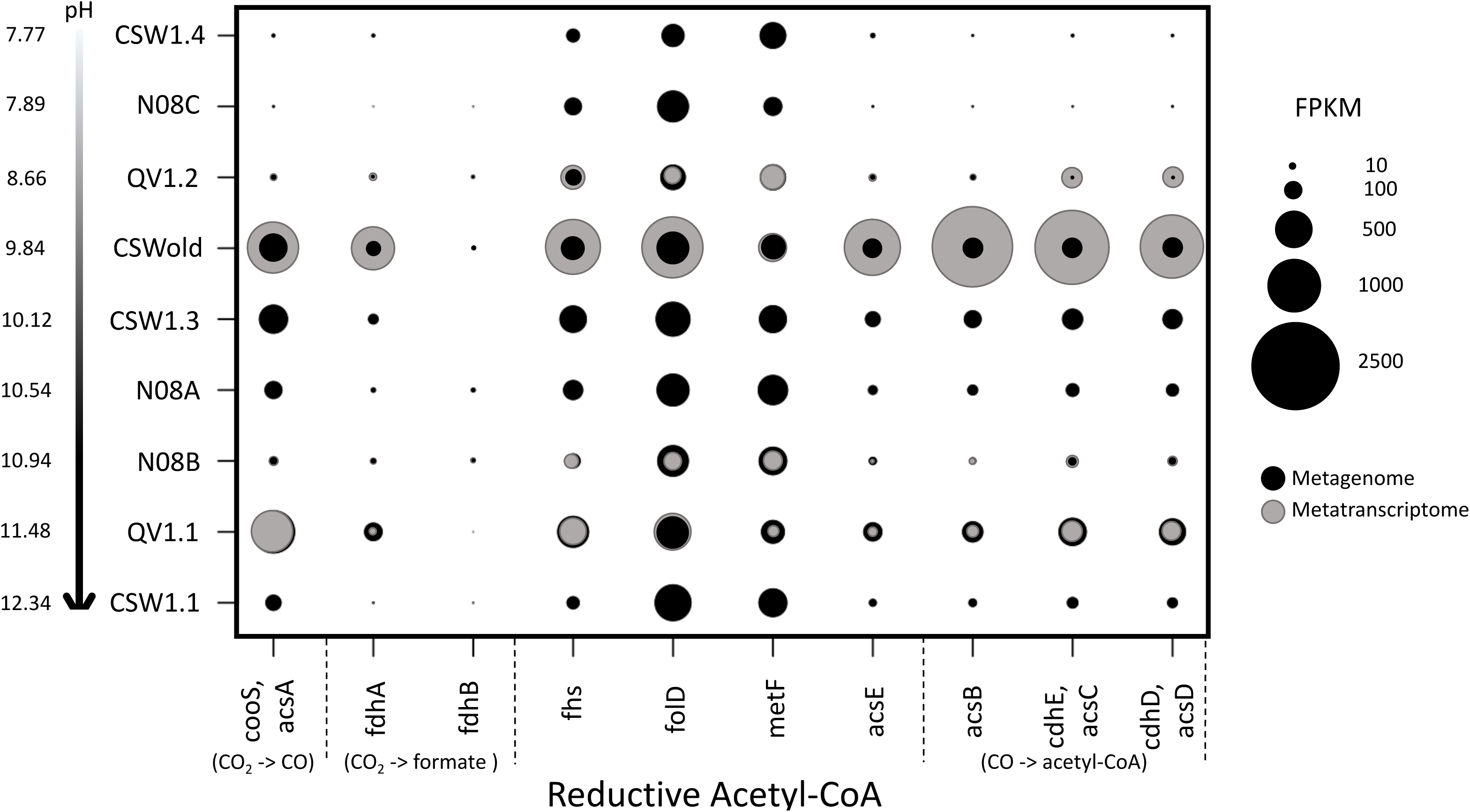
Fragments per kilobase of sequences per million mapped reads (FPKM) of genes in the methane oxidation, tetrahydrofolate (THF), and glutathione-dependent formaldehyde oxidation pathways. Metagenomes are represented in black; metatranscriptomes are represented in gray. The gene for S-(hydroxymethyl)glutathione synthase (gfa) is poorly represented or missing in all wells; this enzyme catalyzes a spontaneous reaction in the formaldehyde oxidation pathway.

### Taxonomy

Taxonomic assignments of the metagenomic contigs using PhyloPythiaS+ were consistent with previous 16S rRNA sequencing data from the CROMO wells (Twing et al., 2017) and showed that the microbial communities in QV1.1 and CSW1.1 (the two most alkaline wells) were highly similar (Supplemental Figure 3). Taxonomic classification of MAG bins, including bin completeness, contamination, and strain heterogeneity, is available in Table 1; a phylogenetic tree comparing the MAGs to the three *Serpentinomonas* isolates is included in Figure 3.

**Table 1.**
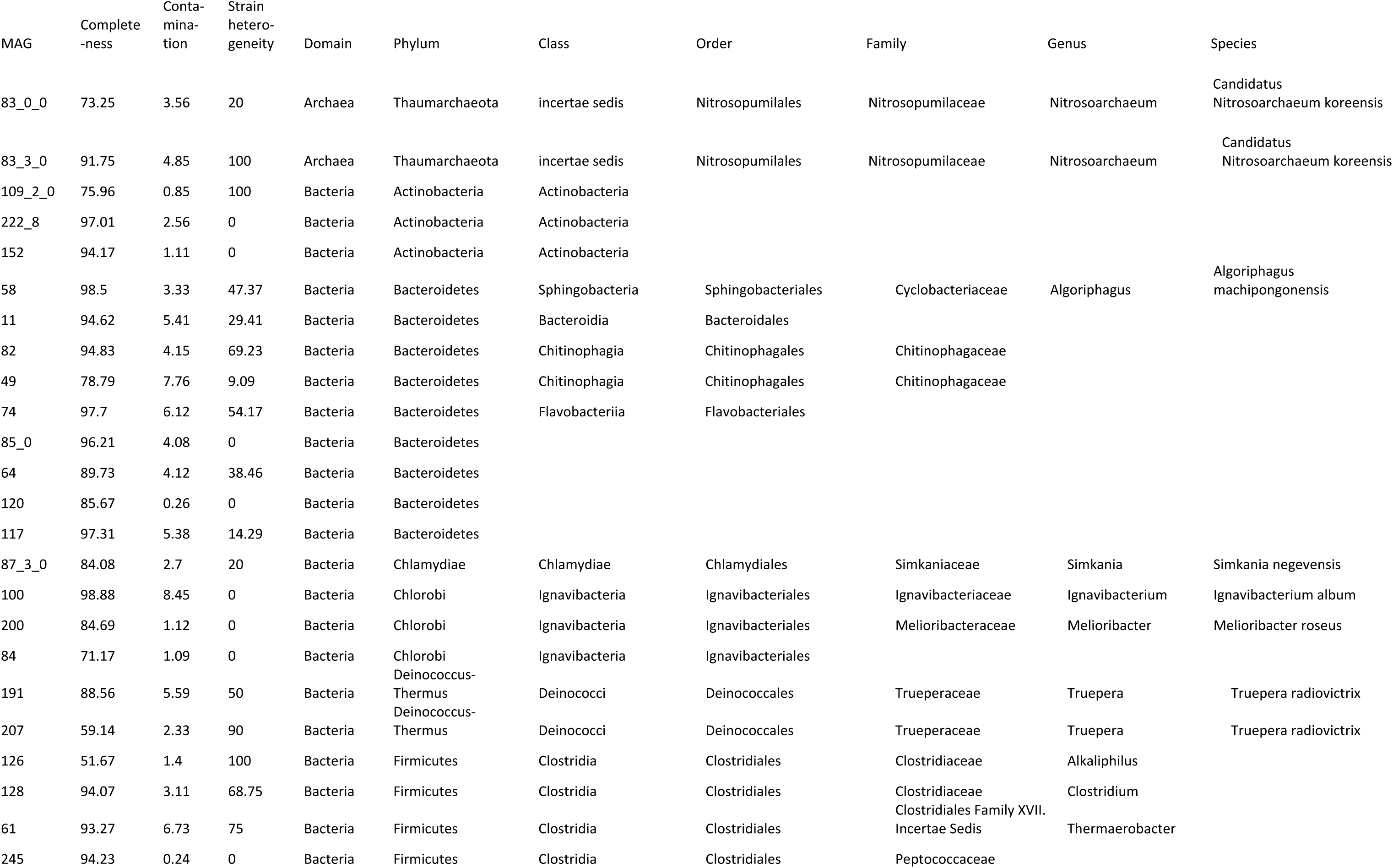
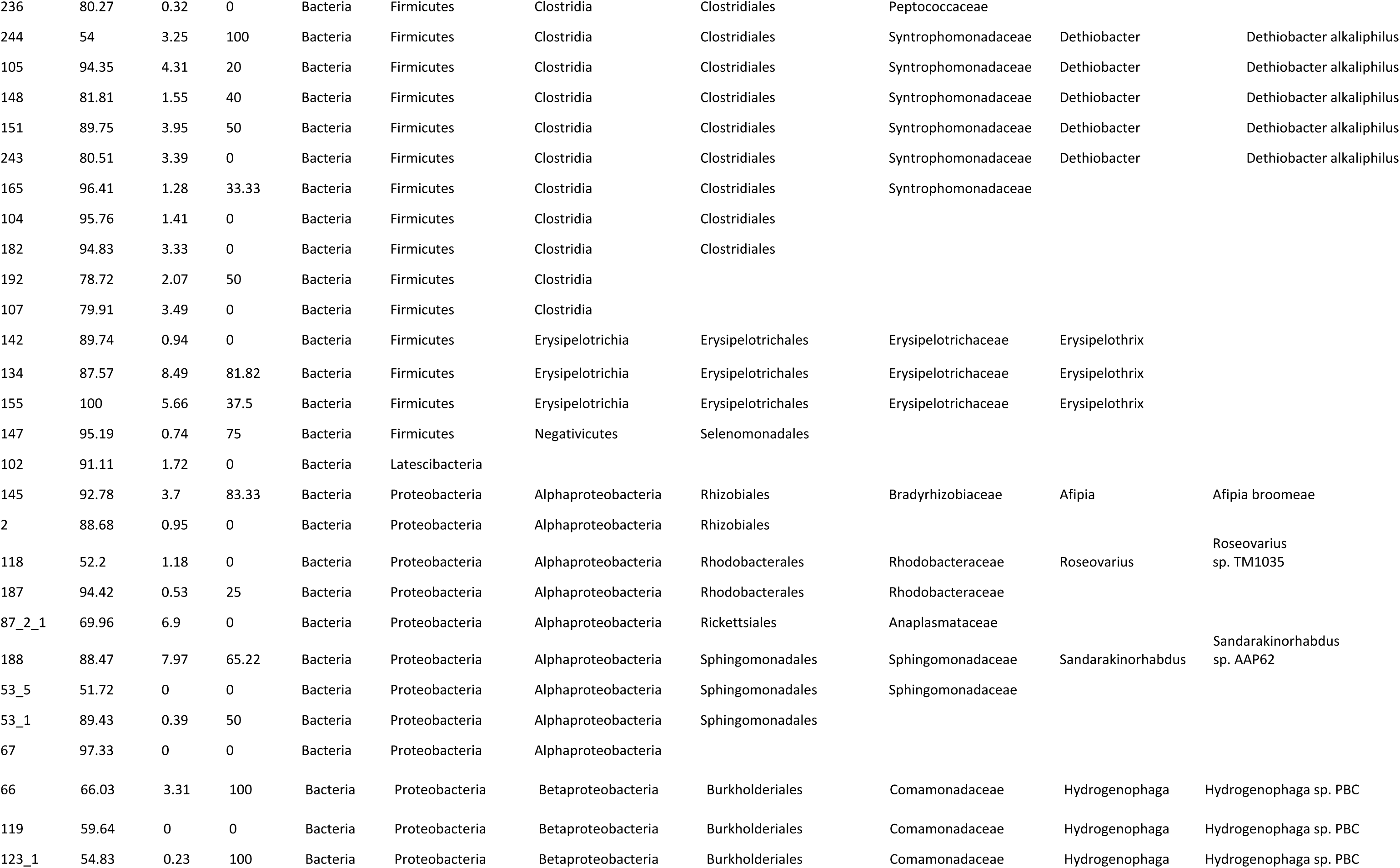
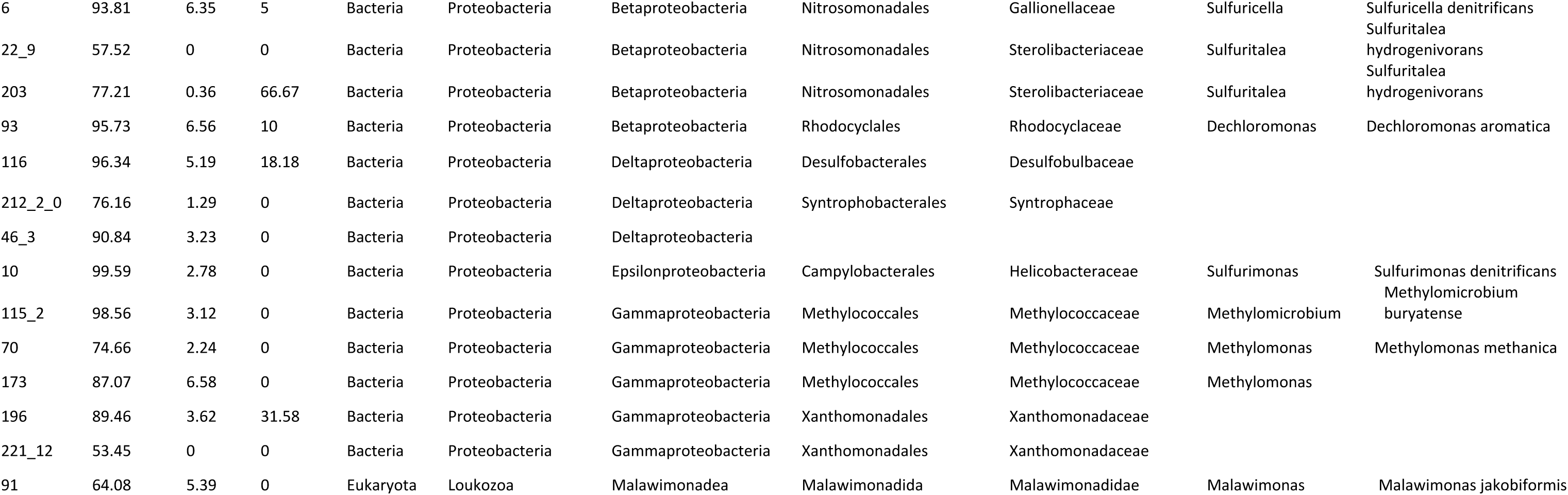
Completeness, contamination, strain heterogeneity, and classification of MAG bins from metagenomes. Bin quality scores assessed using CheckM; taxonomic annotations obtained using GTDB-Tk.

In the metagenomes, *pmoA* and *mxaF* genes from the methane oxidation pathway were detected on contigs classified as *Methylophilales* (*Methylophilaceae*) and *Methylococcales* (*Methylococcaceae*); two MAG bins containing a complete methane oxidation pathway were classified as *Methylococcaceae* (Figure 3; Table 1). Genes from the THF pathway for formaldehyde oxidation (*folD* and *purU*) were associated with *Burkholderiales* (*Comamonadaceae*), *Deinococcales* (*Trueperaceae*), *Hydrogenophilales* (*Hydrogenophilaceae*), *Methylophilales* (*Methylophilaceae*), *Rhizobiales*, and *Rhodocyclales* (*Rhodocyclaceae*); the *frmA* gene from the glutathione pathway for formaldehyde oxidation was detected on contigs identified as *Burkholderiales* (*Comamonadaceae*), *Rhizobiales*, *Rhodobacterales*, and *Rhodocyclales* (*Rhodocyclaceae*). The *folD* gene is also present in the reductive acetyl-CoA pathway for acetogenesis; this gene, as well as *acsB* and *cooS* were associated with *Clostridiales* (*Syntrophomodaceae*) and *Desulfuromodales*. All of the MAG bins containing the reductive acetyl-CoA pathway were classified as *Clostridiales* (Figure 3; Table 1). The large and small subunits of RuBisCO (*rbcL* and *rbcS*) were detected on contigs classified as *Burkholderiales* (*Comamonadaceae*), *Hydrogenophilales* (*Hydrogenophilaceae*), *Rhodocyclales* (*Rhodocyclaceae*), and *Methylophilales* (*Methylophilaceae*). A complete CBB cycle was only identified on one MAG, however, classified as *Rhodocyclales* (*Rhodocyclaceae*). The *aclA* gene from the reverse TCA cycle was associated with *Clostridiales* (*Syntrophomodaceae*) in the metagenomes.

### Metabolomics

The metabolomes of QV1.1 and CSW1.1 displayed many unique observed features across both sample types (intracellular metabolites and extracellular metabolites/dissolved organic carbon [DOC]). In particular, the DOC pools of QV1.1 and CSW1.1 were highly distinct from one another and from intracellular extracts, and these metabolomes contained the greatest number of features unique to the sample (Figure 6). Metabolomes also had more features in common between samples of the same type (e.g., intracellular vs. extracellular) than samples from the same well (Figure 6). Dissolved inorganic carbon concentration in the wells was inversely proportional to pH (Pearson correlation coefficient, r = -0.798), while non-purgeable organic carbon was positively correlated to dissolved oxygen concentration (r = 0.587) (Figure 7). All wells were below the detection limits of the colorimetric formaldehyde test kit (<0.4 ppm).

**Figure 6.**
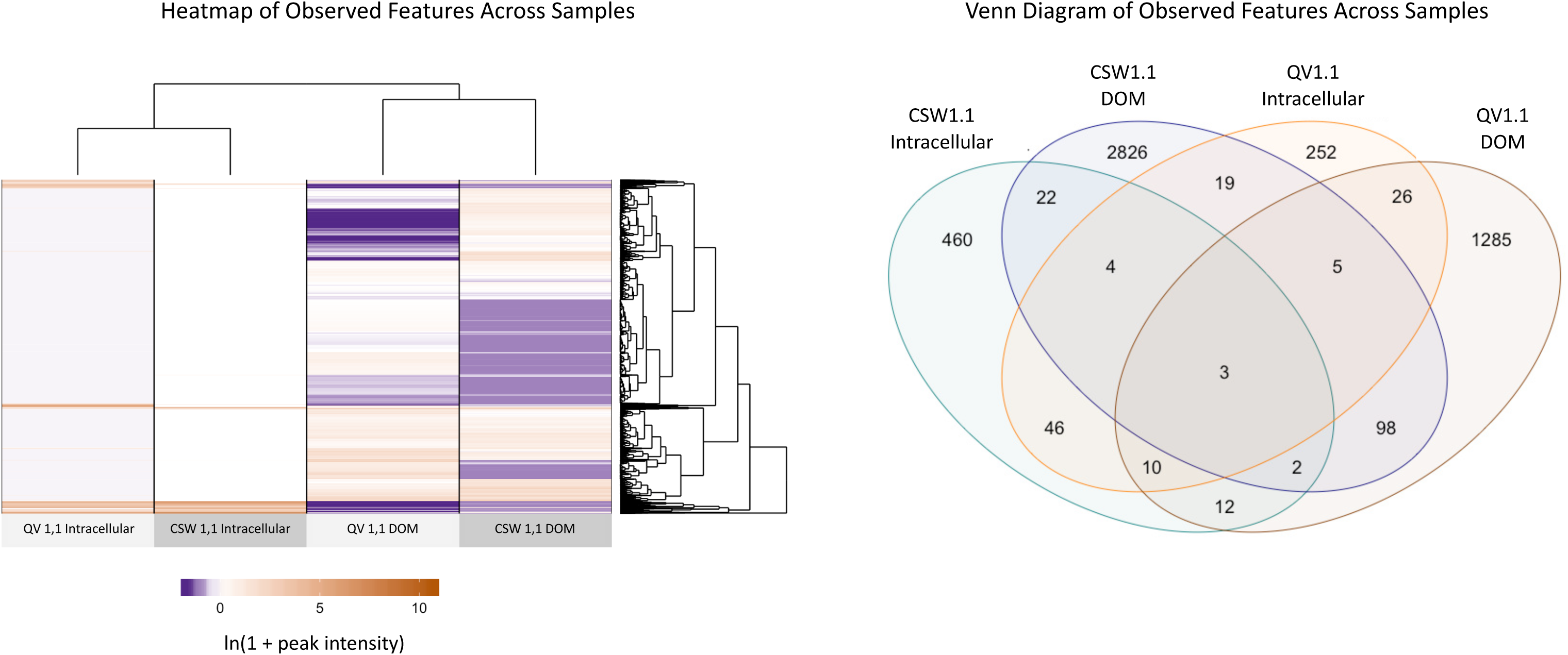
Left: Heatmap of untargeted metabolomics features across all four metabolome samples. The scale represents the ln(1 + peak intensity) (1 is added to avoid cases of ln(0)). Right: Venn diagram of compounds unique to each sample and shared between samples. Observed compounds appear to be generally distinct to each sample. Filtered fluid/dissolved organic carbon samples had the greatest number of unique samples, and CSW1.1 had more unique compounds than QV1.1.

**Figure 7.**
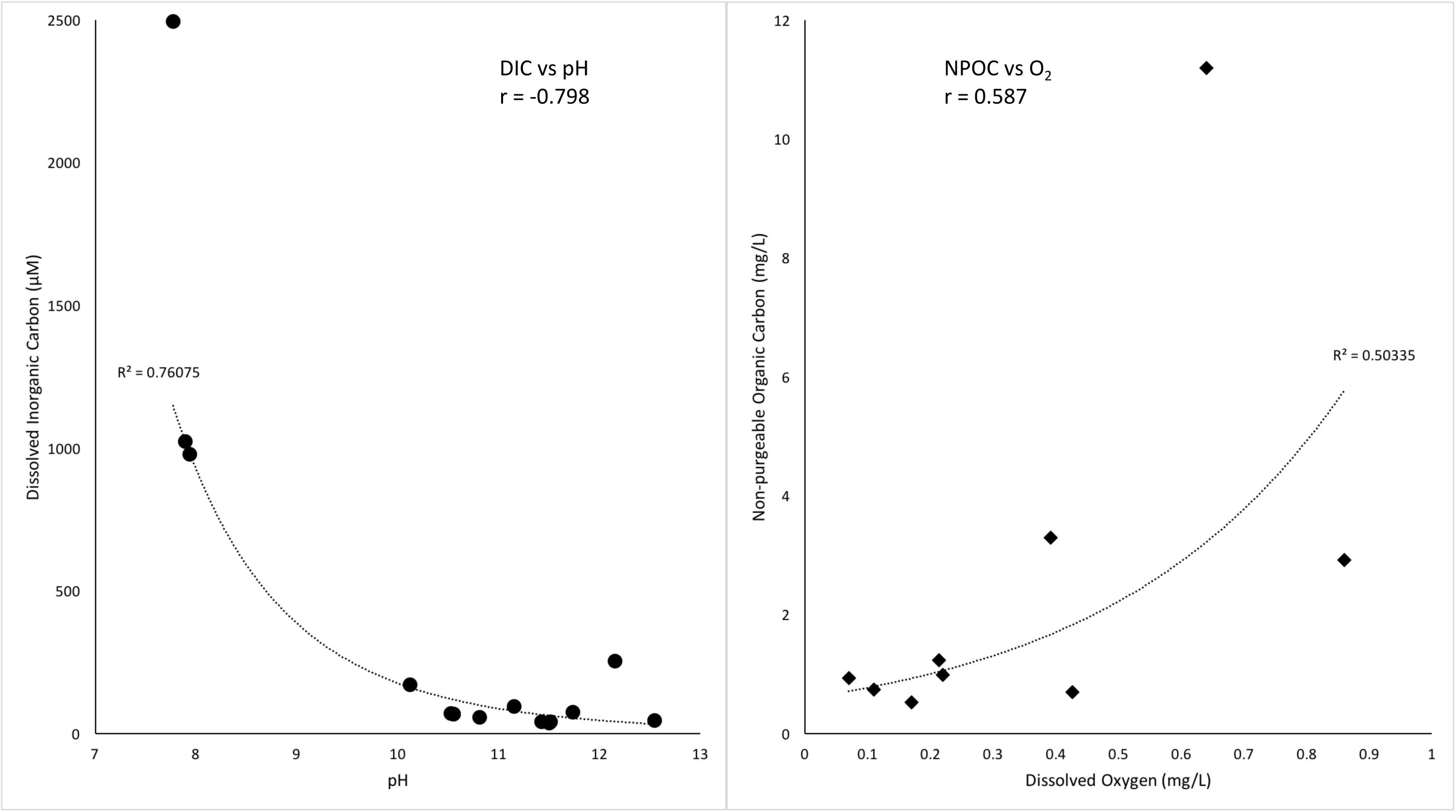
Scatter plots depicting dissolved inorganic carbon (DIC) vs pH (left) and nonpurgeable organic carbon (NPOC) vs dissolved oxygen (DO). The Pearson correlation coefficient (r) is indicated on each graph, as well as the R^2^ of the trendlines.

## Discussion

Previous studies of microbial communities at CROMO have used marker gene analysis, shotgun metagenomics, microcosm experiments, and geochemical measurements to elucidate pathways of biogeochemical cycling in a serpentinite-hosted aquifer (Crespo-Medina et al., 2014; Twing et al., 2017). We expanded upon these efforts by combining metagenomics and metatranscriptomics with an untargeted metabolomics technique to verify genomic expression and metabolic activity. Using this integrated –omics approach, we determined that multiple carbon assimilation pathways are used by the microbial communities in the serpentinite-hosted groundwater at CROMO, including the Calvin-Benson-Bassham cycle, the reverse TCA cycle, acetogenesis via the reductive acetyl-CoA pathway, and methanotrophy.

### Methane Uptake and Formaldehyde Cycling

The addition of methane to microcosms from CROMO has been shown to stimulate microbial growth (Crespo-Medina et al., 2014). In the previous metagenomic study of 4 samples by Twing et al. (2017), no particulate methane monooxygenase (*pmoA*) genes were detected; however, a methanol dehydrogenase (*mxaF*) gene was identified in the CSW1.3 metagenome. Our expanded search through nine metagenomes and four metatranscriptomes succeeding in finding the genes for particulate methane monooxygenase (PMO) and methanol dehydrogenase (MXA) in the CROMO microbial communities, though at low abundance in the higher pH wells (Figure 4), suggesting that the prior perceived absence of these genes could have been the result of a smaller dataset. Two MAG bins contain the complete methane oxidation pathway, and another two contain a partial pathway (Figure 3). No other known methane monooxygenases or methanol dehydrogenases were identified in the metagenomes or metatranscriptomes.

The prevalence of formaldehyde- and formate-assimilation pathways in CROMO reinforces the importance of methanotrophy as a carbon assimilation strategy in these microbial communities. There is a wealth of metagenomic and metatranscriptomic evidence that formaldehyde oxidation to formate and formaldehyde fixation to biomass can be performed by the microbial communities in CROMO (Figure 2). Formaldehyde is an important intermediate in methylotrophic metabolic pathways, and though it is highly toxic, methylotrophic microorganisms have developed a variety of functionally redundant modules for rapidly converting formaldehyde to less toxic compounds (Marx et al., 2005). Formaldehyde can also act as a carbon and electron shuttle between microbial subpopulations, in which it is produced by autotrophic primary producers as a by-product, then taken up by methylotrophic microbes and incorporated into biomass (Moran et al., 2016).

In methanotrophs, formate oxidation to CO_2_ is carried out by the NADP-dependent formate dehydrogenase that converts CO_2_ to formate in the Wood-Ljungdahl pathway, operating in reverse. As previously stated, the genes for this enzyme were not well-represented (and in some cases missing) in the metagenomic data (Figure 5). Formate can also be oxidized to CO_2_ by an NAD-dependent formate dehydrogenase (FDH) as part of the degradation of oxalate; however, the complete suite of genes required to produce this enzyme was likewise poorly-represented or partially missing, particularly in the most alkaline wells (Supplemental Figure 2). This contrasts with subsurface wells in the Samail Ophiolite, where FDH encoding genes were enriched in metagenomes recovered from alkaline and hyperalkaline groundwater (Fones et al., 2019). Therefore, while there is substantial metagenomic and metatranscriptomic evidence for the production of formate from formaldehyde by the microbial communities in CROMO, DNA/RNA evidence for the further oxidation of formate to CO_2_ is lacking.

### Bicarbonate Fixation Strategies in the Metagenomes and Metatranscriptomes

Despite the DIC concentrations being challengingly low for CO_2_ fixation at the high pH of the deepest wells, we found complete Calvin-Benson-Bassham (CBB; Supplemental Figure 1) and reverse tricarboxylic acid (rTCA; Figure 2) cycles in all nine of our metagenomes and all four metatranscriptomes. Sequences encoding the *rbcL* gene of the RuBisCo enzyme were previously detected in metagenomes from four wells (CSW1.1, CSW1.3, QV1.1, and QV1.2) in the earlier metagenomic study by Twing et al. (2017). QV1.1 displays the highest coverage of the rTCA pathway in both its metagenome (607.71±55.83 FPKM) and metatranscriptome (446.86±149.59 FPKM) (Figure 2), even though the concentration of dissolved inorganic carbon (DIC) in QV1.1 is lower than almost any other well (11.48) (conversely, presence/expression of the CBB cycle was roughly equal throughout all the wells [Supplemental Figure 1]). Above pH 11, carbonate is the dominant species of inorganic carbon. *Serpentinomonas* spp., which are among the most abundant taxa in the highest pH wells in CROMO (Twing et al., 2017) require the addition of calcium carbonate (even when bicarbonate is also provided) for growth in culture, and form aggregates on carbonate precipitates (Suzuki et al., 2014). Bicarbonate-fixing microbes may create a localized microenvironment of decreased pH where carbonate may be converted to bicarbonate, or they may have bicarbonate transporters that solubilize carbonate. We did not detect any ATP-binding cassette transporters for bicarbonate in our metagenomic data (Supplemental Table 1). Alternatively, formate may be used as a carbon source in the CBB cycle (Bar-Even, 2016) or in the rTCA cycle (Cotton et al., 2018). Formate could thus be produced from formaldehyde via formaldehyde oxidation and then used in assimilatory/anabolic pathways.

Evidence for the reductive acetyl-CoA pathway was also previously detected in several wells (Twing et al., 2017). In this study, we found a nearly complete homoacetogenic reductive acetyl-CoA pathway in the metagenomic and metatranscriptomic data from all but the two least alkaline wells, except for one gene. The gene that encodes the beta subunit of formate dehydrogenase (*fdhB*), which converts CO_2_ to formate as the first step of the reductive acetyl-CoA pathway, represents <1 FPKM in every metagenome but two (N08A and N08B) (Figure 5), and is only present in the metatranscriptomes of QV1.2 and N08B. Likewise, the genes for FDH are missing from every MAG bin that possesses the reductive acetyl-CoA pathway (Figure 3, inset). Yet, this step of the pathway may be unnecessary for the pathway to proceed if formate is being produced by other means, either through the oxidation of formaldehyde via the glutathione-dependent or tetrahydrofolate pathways, or by abiotic production via Fischer-Tropsch-like reactions. Abiotically-produced formate has been implicated as a possible carbon source in serpentinizing environments such as the Lost City (Lang et al., 2010; Lang et al., 2012; Lang et al., 2018) and the Samail Ophiolite (Fones et al. 2019). While formate-fixing reactions are not common, as formate has a relatively low reactivity compared to CO_2_ or other carboxylic acids (Bar-Even, 2016), formate can be used as the sole carbon source by some microorganisms. In the Samail Ophiolite, rates of assimilation or oxidation of formate were higher than rates of assimilation or dissimilation of bicarbonate or CO regardless of pH (Fones et al., 2019), suggesting that formate may be a preferred source of carbon and/or electrons. Formate concentrations in CROMO are similar to DIC concentrations or lower (Supplemental Table 1; see also Crespo-Medina et al., 2014; Twing et al., 2017), but the addition of formate stimulates microbial growth (Crespo-Medina et al., 2014). We detected two formate transporters in the metagenomic data: *fdhC*, which has been detected in putative formate-metabolizing bacteria in Lost City chimneys (Lang et al., 2018), was detected in two MAG bins (126 and 192, both *Clostridia*), and *focA* was detected in one MAG bin (222_8, *Actinobacter*) and across multiple metagenomes and all four metatranscriptomes (Supplemental Table 1).

As DIC becomes more limiting due to pH, bypassing the conversion of CO_2_ to formate may be an important strategy for carbon assimilation. Provided that sufficient reductant is available (for example, from the oxidation of H_2_), assimilation of CO_2_, CO, or formate can occur via the reductive acetyl-CoA pathway (Takami et al., 2012). The assimilation of formate via the acetogenic reductive acetyl-CoA pathway avoids ATP consumption, supports energy conservation through the use of multiple electron bifurcation mechanisms, and removes the need for an external electron acceptor apart from co-assimilated CO_2_ (Bar-Even, 2016; Cotton et al., 2018). While the *cooS* gene for the conversion of CO_2_ to CO is present (Figure 5), this step could likely be bypassed as well, in wells where the CO concentration is sufficiently high (Supplemental Table 3). Evidence for carbon monoxide uptake (Morrill et al., 2014) and the reductive acetyl-CoA pathway (Suzuki et al., 2017) as important carbon assimilation pathways was previously found in the Tablelands, Canada, and The Cedars, CA, respectively. In the Samail Ophiolite, Oman, it appears that carbon limitation outweighs energy limitation for the autotrophic members of microbial communities at high pH (Fones et al., 2019), making carbon compounds important as both a feedstock for biosynthesis and a source of electrons.

In summary, a putative metabolic pathway for the assimilation of methane into biomass is outlined in Figure 8. Methane oxidation to formaldehyde is carried out by the *Methylococcales* and *Methylophilales*, and then either fixed to biomass via the RuMP and serine pathways, or oxidized to formate by multiple clades via the THF and glutathione-dependent pathways. Formate can then be assimilated into biomass via the reductive acetyl-CoA pathway in the *Clostridiales* or *Desulfuromodales*, as shown, or through the CBB or rTCA cycles (Bar-Even et al., 2016; Cotton et al., 2018).

**Figure 8.**
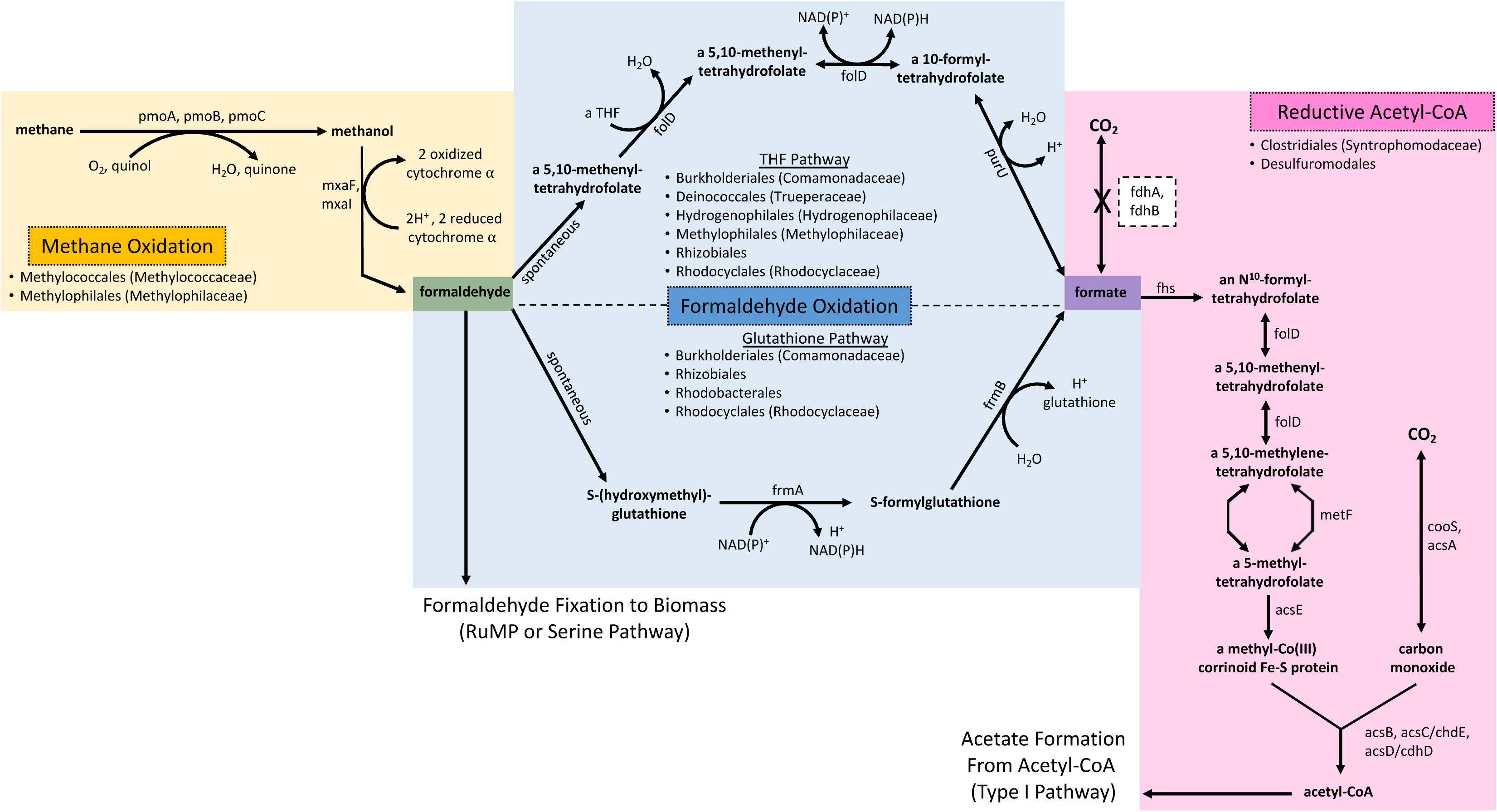
Putative pathway for carbon assimilation in the CROMO wells. Formaldehyde is either converted directly to biomass via the RuMP or serine pathways, or oxidized to formate, which is then fed into the homoacetogenic reductive acetyl-CoA pathway. The pathway for methane conversion to formaldehyde is shown, but appears to be poorly expressed in most of the wells.

### Methanogenesis

While methanogenic archaea have not been detected in 16S rRNA data obtained from CROMO (Twing et al., 2017) and methane isotopologue data suggest a thermogenic source (Wang, 2015), metagenomic and metatranscriptomic evidence suggest the possibility of methanogenesis by multiple pathways in at least some of the wells. However, these pathways do not include the essential backbone of the methanogenesis pathway, including the crucial *mcrA* gene for the production of coenzymeM, required for making methane. The *mcrA* gene was expressed in the metatranscriptomes but other key genes for methane production were absent (Supplemental Tables 1 and 2). The apparent presence of pathways for methanogenesis from organic molecules likely indicate the ability to metabolize these organics (e.g. acetate, methanol, etc.), but whether these compounds are converted to methane by methanogens remains unclear. Biological methanogenesis appears to be occurring in other continental serpentinizing sites, such as the Voltri Massif (Brazelton et al., 2017) and the Cedars (Kohl et al., 2016). In the Tablelands, however, no evidence of biological methane production has been detected (Morrill et al., 2014), and methane production genes in metagenomes recovered from the Samail Ophiolite in Oman are likewise sparse (Fones et al., 2019). Methane cycling in ophiolite-hosted microbial communities therefore seems to vary from site to site, and further work will be required to uncover the geochemical conditions that drive its production and uptake.

### Searching the Metabolome for Intermediates

Intermediates of formaldehyde oxidation and assimilation pathways did not match the masses and retention times of any of the features in our metabolomics data. In negative ion mode, glutathione derivatives should have appeared in the data as deprotonated or double-negatively charged molecules, but we did not detect masses that matched either of those possibilities. This may be due to the relatively low concentrations of DOM (3-13 mg/L) (Fiore et al., 2015), as well as bias introduced by our extraction method (Johnson et al., 2017).

Tetrahydrofolate compounds would likely have been more easily detected in positive ion mode; however, we ran our samples in negative mode in the hopes of capturing the broadest array of metabolites possible with no preconceived notions of what compounds would be present in the data. Metabolites may also fragment or form non-covalent interactions with other compounds upon entering the mass spectrometer, making identification of particular intermediates challenging (Zamboni et al., 2015). Additionally, low cell counts in alkaline groundwater make it difficult to obtain enough biomass during filtration to accurately depict the entire intracellular metabolome of the microbial community. An ideal target of total carbon would be 0.3 mg (Kido Soule et al., 2015). Assuming each cell contains 30 fg of carbon (Fukuda et al., 1998), and with an average cell count of 2 × 10^5^ cells/ml (Twing et al., 2017; unpublished data), 50 liters of water would need to be filtered. Although CSW1.1 and QV1.1 have high fluid outputs compared to most of the other CROMO wells, filtration limitations such as clogging due to carbonate precipitates, keeping the fluid cold to prevent further chemical alteration of the metabolites, and prevention of contamination all precluded our ability to filter such large amounts of water. We were also unable to detect formaldehyde using colorimetric methods in any of the wells. If formaldehyde and its intermediates are cycled rapidly within the cell, it may be difficult to identify and quantify these compounds with the techniques used in this study. Interestingly, while the two wells with the highest pH (QV1.1 and CSW1.1) share highly similar microbial communities (Supplemental Figure 3) with similar metabolic capabilities, the DOC pools observed in these wells appear significantly different in character (Figure 6). Putative annotations of unique metabolite features in the DOC pools include dipeptides, fatty acids, vitamins, phosphorylated metabolites, and secondary metabolites. This observation underscores the need for further characterization of the available carbon in serpentinite-hosted aquifers like CROMO through metabolomics techniques.

### Conclusion

Through an in-depth analysis of metagenomic and metatranscriptomic data taken over several time points, we have determined that methanotrophy, the reductive acetyl-CoA pathway, the reverse TCA cycle, and the Calvin-Benson-Bassham cycle may all be important means of carbon assimilation for microbes living in the Coast Range Ophiolite groundwater. We also identified several mechanisms by which microbes in the hyperalkaline fluids of deep serpentinizing rock may overcome limiting concentrations of DIC, including the use of formate and methane/formaldehyde as carbon sources for homoacetogenesis and the production of biomass. Further optimization of metabolomics techniques for this environment may be of use to track the fate of carbon and delineate between abiotic and biotic processes in serpentinizing systems such as CROMO.

## Acknowledgements

The authors wish to thank McLaughlin Reserve, in particular Paul Aigner and Cathy Koehler, for hosting sampling at CROMO and providing access to the wells, A. Daniel Jones and Anthony Schilmiller for their advice regarding metabolite extraction and mass spectrometry, Elizabeth Kujawinski for her guidance in metabolomics data analysis and interpretation, and Julia McGonigle, Christopher Thornton, and Katrina Twing for assistance with metagenomic and computational analyses.

---

Supplemental Figure 1. Fragments per kilobase of sequences per million mapped reads (FPKM) of genes in the Calvin-Benson-Bassham cycle. Metagenomes are represented in black; metatranscriptomes are represented in gray.

Supplemental Figure 2. Fragments per kilobase of sequences per million mapped reads (FPKM) of genes in the NAD-dependent pathway for formate oxidation to CO_2_. Metagenomes are represented in black; metatranscriptomes are represented in gray.

Supplemental Figure 3. Microbial community composition of QV1.1 and CSW1.1 wells as determined by assigning taxonomy to metagenomic contigs using PhyloPython.

